# Scanning a compressed ordered representation of the future

**DOI:** 10.1101/229617

**Authors:** Zoran Tiganj, Inder Singh, Zahra G. Esfahani, Marc W. Howard

## Abstract

Several authors have suggested a deep symmetry between the psychological processes that underlie our ability to remember the past and make predictions about the future. The judgment of recency (JOR) task measures temporal order judgments for the past by presenting pairs of probe stimuli; participants choose the probe that was presented more recently. We performed a short-term relative JOR task and introduced a novel judgment of imminence (JOI) task to study temporal order judgments for the future. In the JOR task, participants were presented with a sequence of stimuli and asked to choose which of two probe stimuli was presented closer to the present. In the JOI task, participants were trained on a probabilistic sequence. After training, the sequence was interrupted with probe stimuli. Participants were asked to choose which of two probe stimuli was expected to be presented closer to the present. Replicating prior work on JOR, we found that RT results supported a backward self-terminating search model operating on a temporally-organized representation of the past. We also showed that RT distributions are consistent with this model and that the temporally-organized representation is compressed. Critically, results for the JOI task probing expectations of the future were mirror-symmetric to results from memory, suggesting a forward self-terminating search model operating on a temporally-organized representation of the future.

## Introduction

It has been hypothesized that there is a deep symmetry between the ability to remember the past and imagine the future. In episodic memory in particular, the ability to remember episodes from the past has been linked to the ability to simulate the future (Tulving, 1985). Recent findings from neuroimaging studies have found support for this view (e.g., Schacter, Addis, & Buckner, 2007; Bar, 2009; Peer, Salomon, Goldberg, Blanke, & Arzy, 2015). Deficits in the ability to predict the future are known to be comorbid with deficits in episodic memory due to aging (Addis, Wong, & Schacter, 2007) and neurological insults (Hassabis, Kumaran, Vann, & Maguire, 2007). Some authors have even argued that optimal prediction could indeed be the guiding principle underlying the functional organization of the brain (Friston, 2010; Clark, 2013; Bialek, 2012; Palmer, Marre, Berry, & Bialek, 2015) and that there is a deep computational basis for the symmetry between memory and prediction (Gershman, 2017). While many authors have suggested a symmetry between the mechanisms that allow us to access the past and the future, there is a dearth of behavioral paradigms that allow one to directly compare memory access for past and future events under controlled laboratory conditions.

In this paper, we introduce a future-time analog of the relative judgment of recency (JOR) task. By analogy to the way the JOR evaluates participants’ ability to judge the relative time at which past events occurred, this new paradigm tests participants’ ability to judge the imminence of future events over the scale of a few seconds (Figure 1). The classic finding from short-term relative JOR tasks is that correct response time (RT) depends on the lag of the more recent probe but not the lag of the less recent probe (Muter, 1979; Hacker, 1980; Hockley, 1984) as illustrated in Figure 2a. For decades, researchers have argued that this finding is consistent with a self-terminating backward scan along a temporally-organized memory representation (Muter, 1979; Hacker, 1980; Hockley, 1984; McElree & Dosher, 1993). These models implicitly require a memory store that is organized in a way that maps onto the sequential order of experience. Because the scan starts at the present and proceeds backward in time, RTs show a recency effect. The finding that correct RTs do not depend on the lag of the less-recent probe is consistent with a self-terminating search. The data are as if participants stop the scanning process on correct trials once they find the more recent probe. This backward scanning account also naturally explains the finding that incorrect RTs depend on the lag of the (incorrectly) selected probe item under the assumption that errors occur because the scan continues past the correct probe.

**Figure 1.**
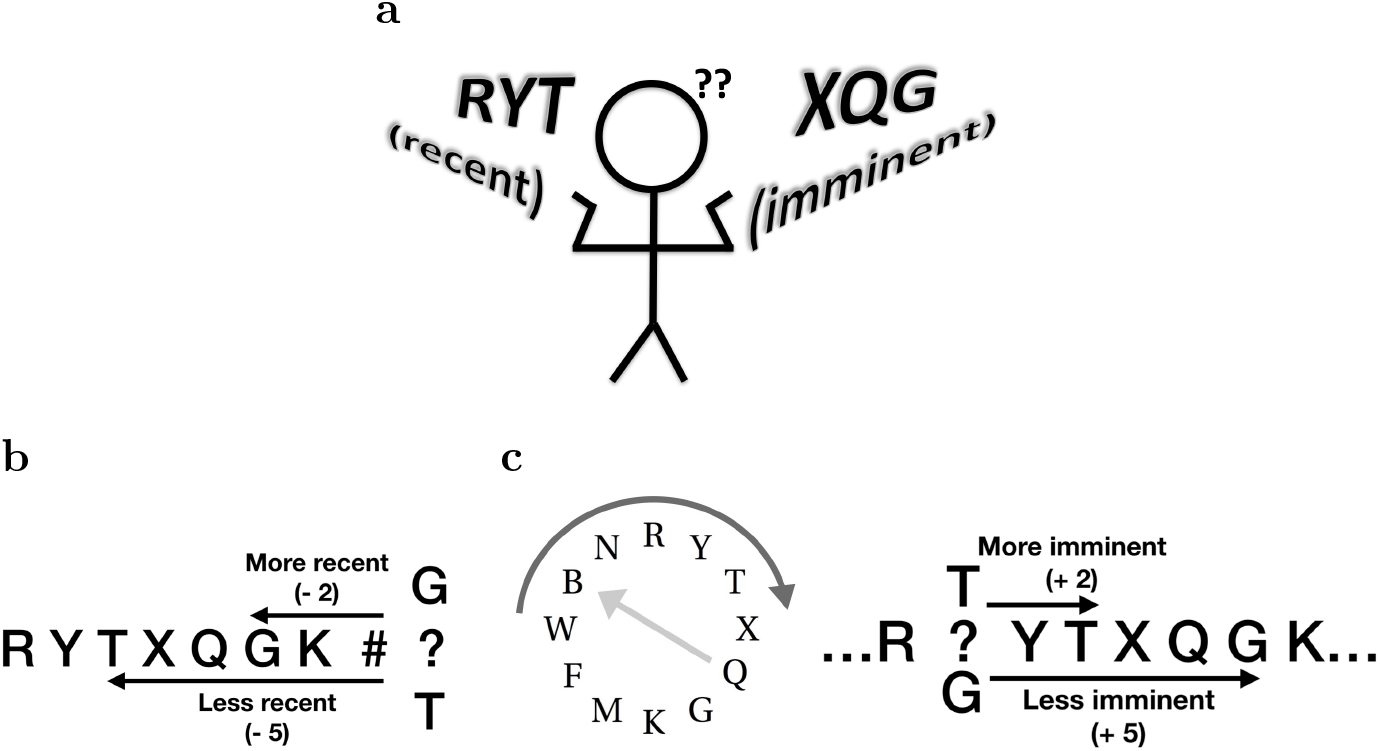
Scanning over ordered representations of the past and the future in judgment of recency (JOR) and judgment of imminence (JOI). **a.** A hypothesis for ordered representations of the past and the future. The memory store for the past is organized in a way that respects the sequential structure of experience. Similarly, the prediction for the future is also organized in a way that respects the sequential nature of the predicted future. The spacing between the letters is foreshortened to suggest compression of the representations for events further from the present. Because events closer to the present are represented with more resolution, the rate of scanning should increase further from the present. **b.** Schematic of the JOR task. Participants are shown a list of letters (such as ryt…) followed by a probe containing two letters from the list (here, g and t). Participants choose the probe item that was experienced more recently. In this example, the probe g is the correct answer. Lag for each probe is defined as the number of steps backward in the list necessary to find the probe. **c.** Schematic of the JOI task. In JOI, participants learn a probabilistic sequence. The sequence usually follows a predictable sequence (clockwise movement along the circle) with occasional random jumps (light-grey arrow). After learning these transition probabilities, the sequence is interrupted by a probe containing two letters (here, g and t). Participants choose the letter that is likely to occur sooner. In this example, the letter t is the correct answer. Lag for each probe is defined as the number of steps forward at which one would expect to find the probe if the sequence continued along its most likely path.

**Figure 2.**
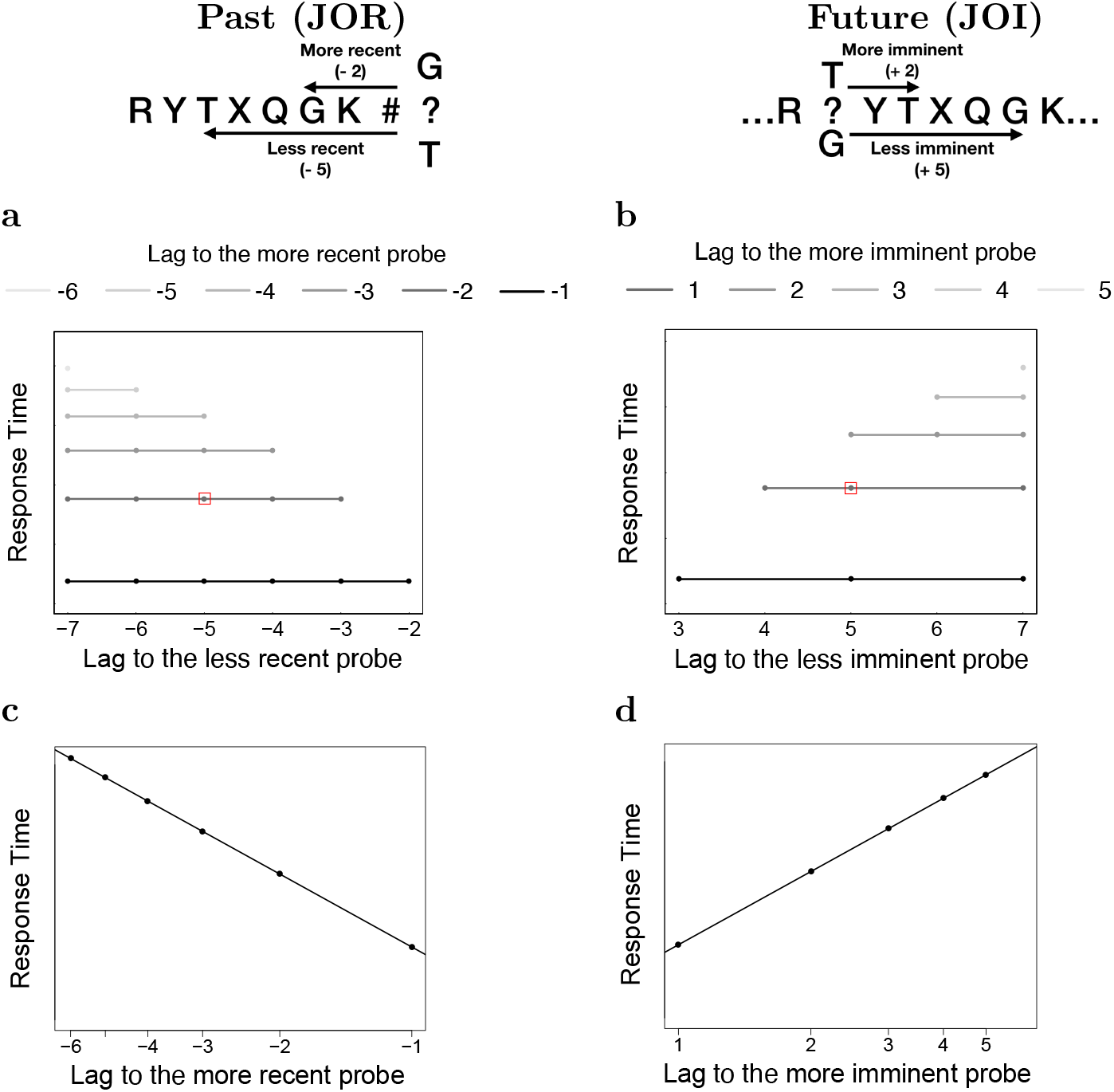
Predictions for a self-terminating scanning model over an ordered compressed representation. **a.** Schematic depiction of RT as a function of the lag of each of the two probes for serial self-terminating scanning model in JOR. Correct RT as predicted from a serial scanning model is shown as a function of the lags of the two probes. The shading represents the lag to the more recent item; the less recent item is plotted on the x-axis. The red square represents the RT corresponding to the probe in the example list. Flat lines for correct RT are a characteristic prediction of serial self-terminating models. This schematic assumes that the memory representation that is being scanned is compressed, such that it takes longer to scan the more recent past than more distant past (notice that the distance between lines representing lag to the more recent probe increases as a function of the lag). The compression is illustrated in plot **c**. If items being scanned are spaced logarithmically along the memory representation then a serial scanning model predicts linear growth of RT as a function of a logarithm of the lag to the more recent item. **b, d.** Anticipated results for JOI if participants use a self-terminating scanning model of a compressed ordered representation of the future. The pattern for correct RTs mirrors the anticipated results for JOR.

A backward scanning model implies that memory has a linear organization that reflects the sequential experience of the list. That is, in order for a memory store to be sequentially “scanned” it must be organized such that adjacent parts of the memory store correspond to adjacent items in the list. This is consistent with longstanding theories of human memory (James, 1890; Brown, Neath, & Chater, 2007; Balsam & Gallistel, 2009; Howard, Shankar, Aue, & Criss, 2015) which argue further that in memory the temporal dimension coding for the past should be increasingly more compressed for a more distant time (Brown et al., 2007; Howard et al., 2015). That is, to account for the finding that memory gets worse for events further in the past, the memory store would devote more resources to recent events and less resources to events further in the past. If the memory store contained a veridical representation of experience, we would expect the time to scan to a probe should be a linear function of its recency—each traversal of a lag in the list should take the same amount of search time. In contrast, the progress made along the list in each moment of serial scanning of a *compressed* memory representation would depend on the degree of compression of that part of the list. Because compression becomes greater for items further in the past, we would expect to see a sublinear increase in RTs with lag (Figure 2c).^1^

Thus far we have detailed predictions of a scanning model on measures of central tendency of RTs. However, scanning models make much stronger predictions. As the “focus of attention” moves backward in time over the memory representation during scanning, it can only acquire information about the region of memory currently in the focus. Consider this process from the perspective of a particular memory from a particular part of the list. Information about that region of memory can only become available after the scan has reached that location. Locations further in the past require additional scanning time before they can start communicating information about the contents of memory in that location. In the language of computational models of RT distributions, the self-terminating scanning model predicts that there should be a recency effect on the non-decision time—a parameter that measures the amount of time that passes before information begins to accumulate (Figure S7, e.g., Ratcliff, 1978). Prior JOR studies have confirmed this prediction by examining RT distributions for probes of different lag (see especially Muter, 1979; Hockley, 1984). Similar conclusions have been reached about the rate of information accumulation using the response signal procedure (McElree & Dosher, 1993).

In order to compare memory for the past with predictions for the future, this study employs a novel judgment of imminence (JOI) paradigm to test predictions of the future. The relative JOR task requires the participant to select the probe item from the past list that was presented closer to the present (Figure 1b). In contrast, the JOI paradigm asks participants to select the future probe item that *is anticipated* closer to the present (Figure 1c). A self-terminating serial search model that scans backward through a compressed record of the past makes a number of predictions for response times—both central tendency and properties of the RT distribution—that are consistent with decades of work in memory research. If memory for the past and prediction of the future rely on the same kind of representation then we would expect that the brain can somehow implement an analogous scanning model over future time. Rather than scanning backward through memory, this model would scan forward through the predicted future to reach an anticipated event. The behavioral predictions of this model would be symmetric to the predictions described above for memory of the past (Figure 2b,d). Observing this pattern of results would suggest that memory and prediction both generate a sequentially ordered representation of events and that we have comparable ability to make judgments about this information in working memory.

## Results

We performed two experiments comparing memory for the past as measured by JOR and prediction of the future as measured by JOI. In Experiment 1 two different groups of participants performed JOR and JOI; in Experiment 2 the two tasks were administered to the same participants in different sessions. Because the results of the two experiments are so similar, we show figures with collapsed data in the main text; figures showing the results of the separate experiments are in the Supplemental Information. Because accuracy from JOR is consistent with previous studies and accuracy for JOI is qualitatively consistent with results from JOR, we discuss accuracy results in the Supplemental information.

To anticipate the results, findings from median correct RTs and careful consideration of RT distributions converged on three primary conclusions. First, the pattern of results in memory and prediction were closely analogous, suggesting symmetry between memory and prediction. Second, the patterns of median RTs from both tasks were as predicted by a serial self-terminating search model over a compressed ordered representation. Finally, a detailed consideration of RT distributions inspired by computational models for evidence accumulation showed strong evidence for scanning in both JOR and JOI. The evidence suggests that participants have access to and can sequentially scan a compressed ordered representation of the future.

### Correct median RT depended only on the lag of the selected probe

Figures 3a and b show median correct RT as a function of the lag of both of the probe stimuli collapsed over the two experiments. Results for JOR, evaluating memory for the past, are shown in Figure 3a; results for JOI, evaluating prediction of the future, are shown in Figure 3b. The results for the separate experiments are shown in Figure S2. Visual inspection of this figure shows results consistent with a serial self-terminating search model. First, there was a large proximity effect such that when the correct probe was closer to the present, RTs were faster. This can be seen by noting that the darker lines are lower than the lighter lines. Second, each of the lines in Figures 3a and b appears to be flat, indicating that the lag to the less recent probe did not affect response time.

**Figure 3.**
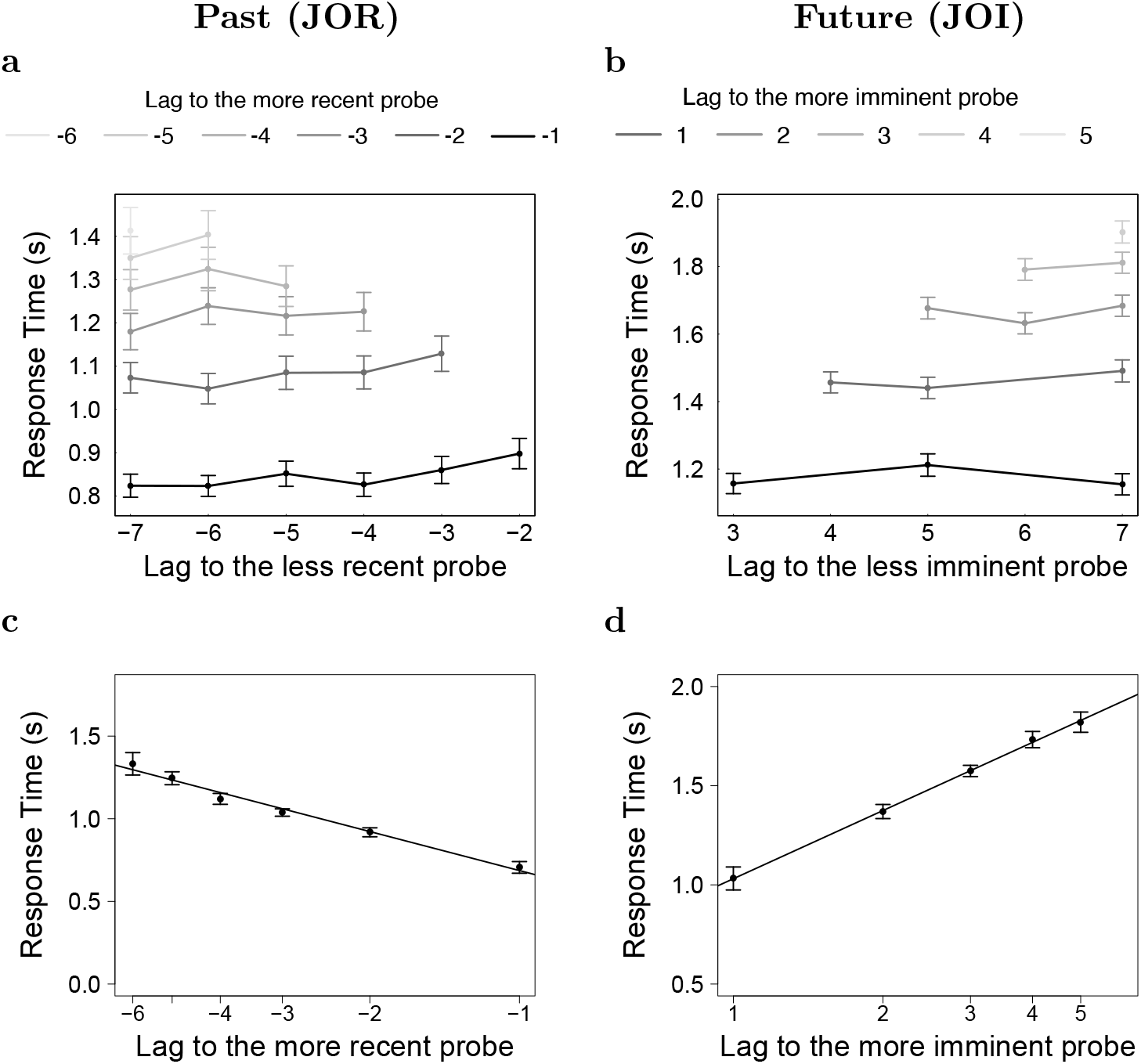
RT depends on the lag of the more recent/imminent probe but not on the lag of the less recent/imminent probe. Median RT varies sub-linearly with lag. Error bars represent the 95% confidence interval of the mean across participants. **a.** In JOR, median RT for correct responses depends strongly on the lag of the more recent probe (indicated by different colors) but not on the lag of the less recent probe (x-axis). **b.** In JOI, median RT for correct responses depends strongly on the lag of the more imminent probe (indicated by different colors) but not on the lag of the less imminent probe (x-axis). **c.** In JOR, median RT varies sub-linearly with lag of the more recent item. **d.** In JOI, the median RT varies sub-linearly with lag of the more imminent item.

#### JOR: Memory for the past

Statistical analyses confirmed these impressions for the JOR data from each experiment taken separately. In Experiment 1, a linear mixed effects analysis allowing for independent intercepts for each participant showed a significant effect of the lag to the more recent probe, .124±.006 s, *t*(478) = 21.6, *p* < 0.001. In Experiment 2, there was also a significant effect of the lag to the more recent probe, .107 ± .009 s, *t*(153) = 11.95, *p* < 0.001. To assess whether there was an effect of the less recent probe, we calculated the slopes of each of the lines in Figure 3a separately for each participant and performed a Bayesian t-test (Rouder, Speckman, Sun, Morey, & Iverson, 2009) on the slopes. In Experiment 1, this analysis showed “substantial evidence” (Wetzels & Wagenmakers, 2012; Kass & Raftery, 1995; Jeffreys, 1998) favoring the hypothesis that the slopes are not different from zero (JZS Bayes Factor = 3.3). Experiment 2 also provided “strong” evidence for the hypothesis that the slopes are not different from zero (JZS Bayes Factor = 10.91). These results replicate prior studies of JOR, but extend them by establishing positive evidence for the null distance effect using the Bayesian t-test.

#### JOI: Predicting the future

Statistical analyses led to the same conclusions for JOI. In Experiment 1, a linear mixed effects analysis allowing for independent intercepts for each participant showed a significant effect of the lag to the more imminent probe, .195 ± .009 s, *t*(227) = 24.1, *p* < 0.001. Results from Experiment 2 also showed an effect of lag to the more imminent probe, .191 ± .016 s, *t*(91) = 11.6, *p* < 0.001. A Bayesian t-test on the slopes of the JOI data (Figure 3b) from both Experiments showed “substantial evidence” favoring the hypothesis that the slopes are not different from zero (Experiment 1, JZS Bayes Factor = 9.17; Experiment 2, 4.76).

### Correct RT varied sub-linearly with lag of the selected probe, consistent with scanning over a compressed representation

The separation between the nearly-horizontal lines in Figures 3a and b appears to decrease as the more recent probe is chosen to be further and further in the past. Figures 3c and d display median correct RT collapsed over the two experiments as a function of the lag to the *more* recent probe on a logarithmic scale. Results from each of the two experiments are shown separately in Figure S3. In both JOR and JOI, correct responses are faster when the selected probe is closer to the present. This manifests as a recency effect in JOR, which can be seen as a line that decreases from left to right in Figure 3c. JOI also shows a proximity effect, which manifests as the line increasing from left to right in Figure 3d. Because the *x*-axes are compressed, to the extent RTs appear as a straight line when plotted in this way, one can infer that the increase in RT is sublinear. That is, if there was a linear relationship between RT and lag, when plotted on a compressed *x*-axis the line would appear as a curve accelerating upwards towards the left of the Figure 3c for JOR and upwards to the right of Figure 3d for JOI.

Statistical analyses confirmed the visual impression that the increase in RT was sub-linear for both JOR and JOI. To assess this we compared a regression model of median RT onto lag to a regression model onto the base-2 logarithm of the absolute value of lag. To the extent the model using base-2 logarithm as a regressor provides a better fit, we can conclude that the increase in RT is sublinear.^2^

#### JOR: Memory for the past

For Experiment 1, the model using log |lag| fit better than the model using |lag|, Δ*LL* = 2.0, implying that the model using the logarithm is about 7 times more likely. Similar results were found for Experiment 2, Δ*LL* = 2.6, implying that the model using log |lag| is about 13 times more likely than the model using |lag|. To quantify the relationship between correct RT and log |lag|, we performed a linear mixed effects analysis allowing for independent intercepts for each participant. In Experiment 1, this analysis showed that every doubling of |lag| increased RT by .24 ± .01 s, *t*(478) = 21.74, *p* < 0.001. Similar results were observed in Experiment 2, where each doubling of |lag| increased RT by .21 ± .02 s, *t*(153) = 12.27, *p* < 0.001. The finding of sublinearity in JOR is consistent with prior studies (Hacker, 1980; Hockley, 1984). However to our knowledge it had not previously been statistically evaluated.

#### JOI: Predicting the future

In Experiment 1 the model using log lag as a regressor fit better, Δ*LL* = 16.3, implying the model using log lag is about 10^7^ times more likely. In Experiment 2, similar results were observed, with Δ*LL* = 5.8, suggesting the model using log lag is about 350 times more likely. A linear mixed effects analysis allowing for independent intercepts for each participant showed that each doubling of lag increased the RT by .34± .01 s, *t*(227) = 26.5, *p* < 0.001 in Experiment 1. In Experiment 2, each doubling of lag led to an increase of .34 ± .03 s, *t*(91) = 12.78, *p* < 0.001.

### Evidence for scanning from RT distributions

The preceding results showed that both memory and prediction showed a proximity effect in the time to correctly select a probe. This result is consistent with a self-terminating serial scanning model operating on a sequentially-organized compressed representation. The scanning model makes a much stronger quantitative prediction that goes beyond measures of central tendency in RT. Because information about a probe item should only become available when attention reaches the part of memory where the probe is found, this model predicts that the entire RT distribution for correct responses should be shifted as a function of the distance that must be scanned to find the probe.

Our hypothesis requires us to evaluate whether the non-decision time changes with proximity. The additional supplemental material shows cumulative RT distributions separately for each participant in this study. To evaluate our hypothesis in the simplest way possible, we use a minimal non-parametric estimate of non-decision time. Consider how quantile plots as a function of log |lag| would look if the proximity effect was carried by a change in non-decision time or if it was due only to a change in drift rate (Figure S7). Because a change in non-decision time would move the entire RT distribution to the right, we would expect to see parallel curves as a function of log |lag| (Figure S8a right hand side). The particular shape of the RT distribution would control the spacing between the lines, but if the effect of log |lag| is solely *via* the non-decision time, the lines connecting any particular quantile across lags should have the same slope. In contrast, if the effect of log |lag| was carried *via* drift rate, then the rate of information accumulation should decrease monotonically with the quantile chosen and result in an asymptotic slope of zero at very small quantiles (Figure S8a left hand side). Of course it is possible that recency affects both the non-decision time and rate of information accumulation. In this case, we would expect to see a combination of effects; the characteristic signature of a change in non-decision time is the non-zero slope as the quantile becomes very small.

Figure 4 shows RT quantiles as a function of log |lag| for both JOR and JOI collapsed over experiments (results for the two experiments separately can be found in Supplemental Figure S10). For JOR, the curves appear to increase from right to left for every choice of quantile, including the lowest quantile at the bottom of the plot. In addition, there also appears to be a systematic change in the slope of the line, consistent with a change in the rate of information accumulation as a function of recency. The results for JOI are particularly striking, appearing to show parallel straight lines as a function of log lag. The curves not only increase from left to right at small quantiles (at the bottom of the plot), but increase by the same amount regardless of quantile. This suggests that in JOI, the effect of lag on RT was carried almost exclusively by a change in the non-decision time that was a function of log lag.

**Figure 4.**
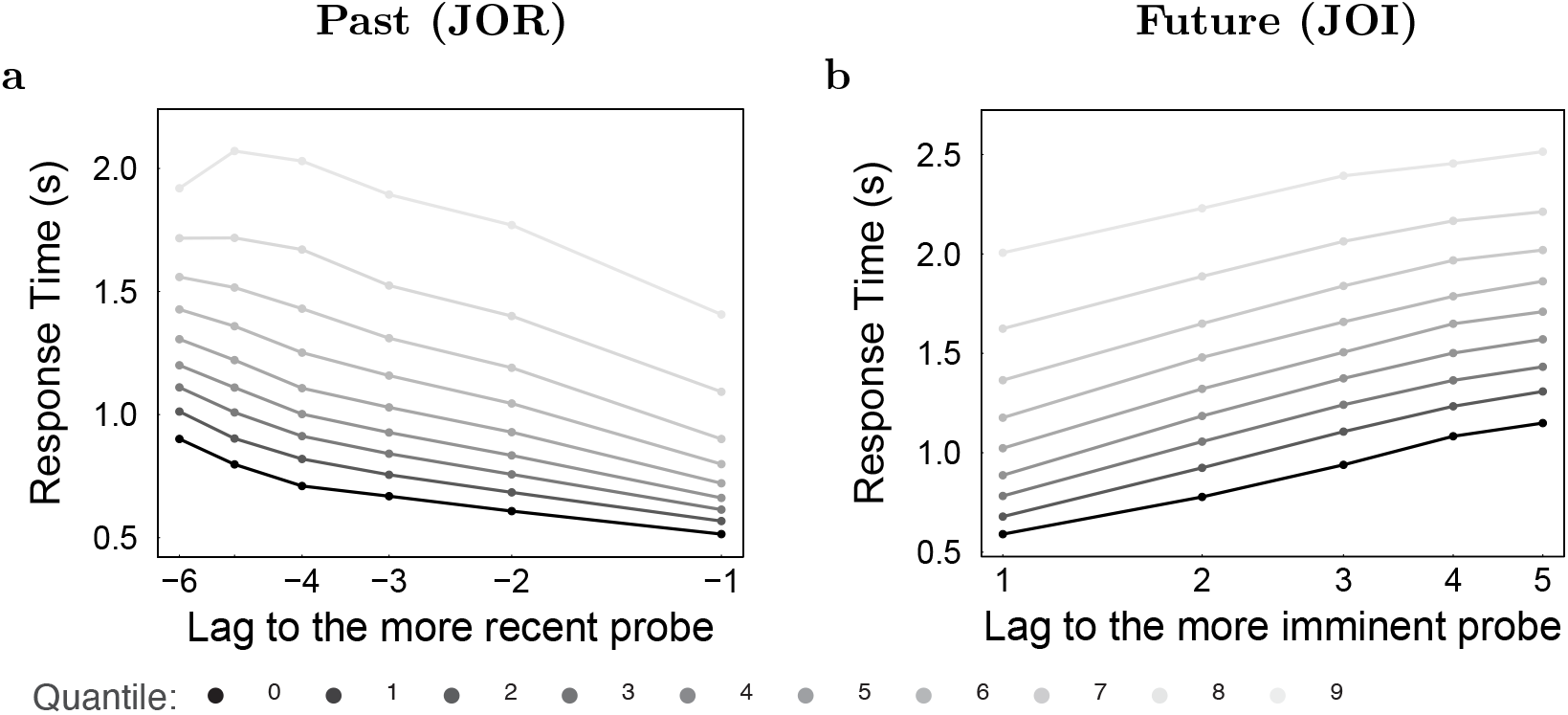
Properties of RT distributions were consistent with the scanning hypothesis in both JOR and JOI. Response time as a function of lag for every decile across all the participants (together, Experiment 1 and Experiment 2). Visual inspection suggests that the results resemble the “non-decision time only” (consistent with serial search models) hypothesis over the “drift rate only” hypothesis (consistent with parallel search models). This is supported by a statistical analysis shown in the main text. See Figure S8 for predictions of the “drift rate only” and “non-decision time only” hypotheses.

To quantify this pattern of results, we define a “Lag Modulation Factor” as the slope of the regression of RT at a particular quantile onto log |lag| (Figure S8b). If the proximity effect changes the non-decision time, as predicted by the scanning model, we would expect the Lag Modulation Factor to be constant and non-zero as quantile goes to zero. We can summarize this prediction by taking the slope and intercept of the line relating Lag Modulation Factor to quantile in Figure S9a. If the scanning hypothesis holds, then across participants we would expect to see non-zero intercepts but slopes around zero (insert in Figure S8b right). In contrast, if proximity changed the drift rate, the change in RT should be primarily due to changes in the tails of the RT distribution. In that case, the effect of lag should decrease as one considers lower and lower quantiles. If lag affects the rate of information accumulation, then the Lag Modulation Factor should decrease continuously as the quantile approaches zero, leading to very different predictions (insert in Figure S8b left).

Figure S9a shows the Lag Modulation Factor as a function of quantile for each participant in Experiments 1 and 2. The lines appear to be flat with a non-zero intercept as the quantile decreases, as predicted by the scanning hypothesis. Once again, the results from JOR and JOI were closely analogous. The scatterplot in Figure S9b shows each participants’ slope and intercept for each experiment and each task. Visually, these results clearly support the scanning hypothesis at the group level. Statistically, in JOR Experiment 1 and Experiment 2 a Bayesian t-test showed “decisive evidence” favoring the hypothesis that the intercepts were different from 0 (JZS Bayes Factor > 100 in both experiments), as predicted by a change in the non-decision time with recency. In addition, a Bayesian t-test also showed “strong evidence” for the hypothesis that the slopes were different from 0 (JZS Bayes Factor = 17.0 in Experiment 1 and JZS Bayes Factor = 11.9 in Experiment 2), suggesting that there was also an effect on the rate of information accumulation. In JOI statistical results were even stronger in support of the scanning hypothesis. As in JOR, for intercepts in Experiment 1 and Experiment 2 a Bayesian t-test showed “decisive evidence” favoring the hypothesis that they were different from 0 (JZS Bayes Factor > 100 in both experiments), suggesting that there was a systematic change in the non-decision time with proximity. In addition, a Bayesian t-test also showed “strong evidence” that the slope was not different from 0 (JZS Bayes Factor = 5.9) in Experiment 1 and “substantial evidence” (JZS Bayes Factor = 2.0) in Experiment 2. This suggests that there was also an effect of proximity on the rate of information accumulation.

## Discussion

The goal of this study was to develop a paradigm to investigate predictions of the future that is analogous to the relative JOR task used to study memory for the past. This study found that judgments of the imminence of future events have the same properties as judgments of the recency of past events. In both cases, the results are consistent with scanning models that posit that information is stored along an organized representation that can be sequentially accessed (Murdock, 1974). Because we did not separately manipulate time and ordinal position in either experiment, the present study does not disambiguate temporal and ordinal representations (Hintzman, 2004). In either case, the scanning hypothesis requires some form of sequential organization to memory and prediction. The major difference between memory and prediction is the direction of the scan—towards the past in the case of memory but towards the future in the case of prediction. In both cases, however, the scan extends away from the present, suggesting a symmetry between memory and prediction.

Critically, scanning of both the past and the future suggests that the temporal representation is compressed, with decreasing accuracy for events further from the present. This compression is reflected in the sublinear increase of correct RT with distance from the present (Figure 3c,d). In both cases we can certainly reject the hypothesis that RT increases linearly. The results are consistent with a number of forms of compression and we cannot make a strong conclusion about the precise quantitative form of the compression. However, the results are at least consistent with a logarithmic compression. Logarithmic compression is implied by the Weber-Fechner law and implements a scale-invariant temporal support, as predicted by quantitative models of memory (Brown et al., 2007; Howard et al., 2015). In vision, it is well known that retinal coordinates are approximately logarithmically-compressed (Daniel & Whitteridge, 1961; Hubel & Wiesel, 1974; Schwartz, 1977). Logarithmic compression of RTs as a function of proximity would suggest a deep connection between judgments of lag and perception, suggesting that time in memory behaves something like the physical sense of vision (Howard, 2018).

The similarity of the results for memory and prediction were observed despite numerous procedural differences between the tasks. First, the lists were presented at a much faster rate in JOR than in JOI. In JOR the list was presented quite quickly, with one item about every 180 ms, such that the entire list was complete in less than 3 s. In contrast, in JOI, the list was experienced much more slowly: a letter was presented every 1.4 s such that a single presentation of the list took about 18 s. Second, the participants in JOI had only a single list to remember throughout the session. In contrast, in JOR participants learned many lists using the same stimuli over the course of a session. In light of these procedural differences between the JOR task and the JOI task, the similarity of the core results is even more striking. Despite these differences the scanning rates across tasks were comparable, with about 220 ms per doubling of |lag| in JOR and about 340 ms per doubling in JOI.

Previous studies (Chan, Ross, Earle, & Caplan, 2009) have reported that subtle differences in instructions given to participants can bias memory search. Chan et al. (2009) found that when participants are instructed to judge which of the two probe items was presented earlier they performed a forward search (RT depended primarily on the lag of a less recent item). When Chan et al. (2009) changed the wording of the instruction such that participants were asked to judge which probe item was presented later they performed a backward search, just as in the present study. An analogous manipulation is not possible in the JOI task used here because participants were trained on a continuous sequence. Future studies could explore the importance of instructions in JOI by changing the experimental paradigm such that participants are trained on lists of items with beginning and end, or perhaps lists containing distinctive items that function like landmarks. Scanning along the timeline of the future suggests that humans are able to deploy attention to different parts of the mental timeline. Evidence for this was shown in Denison, Heeger, and Carrasco (2017) where participants were able to direct attention to a cued point in time. Furthermore, a study conducted in parallel to this one indeed found evidence that, when instructed, participants were able to deploy attention to a several seconds wide temporal window located several seconds into the future (Babcock, Howard, & McGuire, 2019). In a recent study, (Grabenhorst, Michalareas, Maloney, & Poeppel, 2019) described the interplay of the hazard rate of event probability and the uncertainty in time estimation as two main parameters that impact brain’s ability to estimate the future. While we did not manipulate the event probability, the sublinear increase of RT with distance from the present observed in JOI is consistent with the increase of uncertainty in time estimation.

### Computational approaches for constructing a compressed ordered representation of the future

Reinforcement learning models have been extremely influential in the cognitive neuroscience of prediction (Schultz, Dayan, & Montague, 1997; Daw & Dayan, 2014). In this view, the brain can predict the future using either a model-free or a model-based system; these two systems have different properties and can be brought to bear under appropriate circumstances (e.g., Daw, Niv, & Dayan, 2005; Russek, Momennejad, Botvinick, Gershman, & Daw, 2017; Momennejad et al., 2017). The results of the experiments in this paper are difficult to reconcile with either the model-free or model-based systems as currently understood.

Model-free learning efficiently extracts predicted outcomes that follow from a stimulus. A major benefit of model-free learning is that knowledge of outcomes is cached so that they can be rapidly accessed, avoiding the relatively costly computation associated with the model-based system. For instance, the successor representation (Dayan, 1993) is a computational method that enables rapid learning of exponentially-weighted future outcomes that follow a particular stimulus. The successor representation has many properties that have made it attractive to cognitive neuroscientists (e.g., Stachenfeld, Botvinick, & Gershman, 2017; Momennejad et al., 2017). Because the successor representation more strongly weights events that are likely to happen closer in time to the present, it would be straightforward to predict a lag effect in JOI using the successor representation to generate a strength for each probe. Moreover, this effect of lag would be sublinear because as the successor representation weights future outcomes in an exponentially-discounted manner. This is at least roughly consistent with the results in Figure 3d. However, because the strengths of both probes are cached, there is no reason to expect that there would not be a strong interaction between the two probes and a robust distance effect in RT. This is in sharp contrast to the lack of a distance effect on correct RT (horizontal lines in Fig. 3b) and the account of these results based on scanning. Similarly, there is no reason to expect that the time it takes to retrieve a cached strength depends on proximity, so it would be difficult to reconcile the RT quantile plots in JOI (Figure 4) with the output of the successor representation.

In contrast, a model-based account is perfectly consistent with a scanning hypothesis. The model-based system is believed to store the transition probabilities between stimuli (Daw et al., 2005). Predictions at successive time points can be simulated by repeatedly applying the matrix of transition probabilities to the current state. At each time step, one could compare the predicted future to the probes and stop when a match is found. Although the model-based system is well-positioned to build a scanning model, it is not clear why the growth in RTs would be sublinear. If one accepts the quantitative argument that the growth of RTs is logarithmic, which implies that the rate of applying the transition matrix must somehow increase exponentially in order to account for the data.

We have seen that the behavioral findings from JOI are consistent with serial self-terminating search of a compressed ordered representation of the future. Computational behavioral modeling (Howard et al., 2015; Tiganj, Cruzado, & Howard, 2019) has shown that RTs in JOR can be modeled using a logarithmically compressed ordered representation of the past. The representation of the past used in that model corresponds closely to time cells (Pastalkova, Itskov, Amarasingham, & Buzsaki, 2008; MacDonald, Lepage, Eden, & Eichenbaum, 2011; Eichenbaum, 2014), populations of neurons that have been observed in a variety of brain regions, including prefrontal cortex (Tiganj, Kim, Jung, & Howard, 2015; Tiganj, Cromer, Roy, Miller, & Howard, 2018; Cruzado, Tiganj, Brincat, Miller, & Howard, 2019) suggesting an involvement in working memory. Coupled with attentional mechanisms from visual search and evidence accumulation circuits, that model is composed entirely of components that can be, in isolation at least, taken literally as computations the brain could perform.

Can a similar strategy be used to construct a neurally plausible model of the JOI task, scanning the future? Because the behavioral results from JOI are so consistent with the behavioral results from JOR, the primary requirement for a neurocognitive model of JOI is a neurally-plausible compressed representation of the future. Computational work has developed several neurophysiologically-reasonable hypotheses for constructing a compressed ordered representation of the future (Shankar, Singh, & Howard, 2016; Tiganj, Gershman, Sederberg, & Howard, 2019; Momennejad & Howard, 2018; Tano, Dayan, & Pouget, 2020). A compressed record of the past—time cells—can be used to generate a compressed estimate of future events (Shankar et al., 2016; Tiganj, Gershman, et al., 2019). The Shankar et al. (2016) model suggests a connection between the prediction of the future and theta oscillations, a 4-8 Hz rhythmicity that can be observed in humans (Herweg, Solomon, & Kahana, 2020). The Tiganj, Gershman, et al. (2019) paper utilizes simple Hebbian plasticity between time cells and cells coding for present stimuli. Momennejad and Howard (2018) noted that a compressed ordered estimate of future events could be generated from an ensemble of successor representations with different planning horizons and noted the correspondence with neurophysiological findings (Ferbinteanu & Shapiro, 2003; Gauthier & Tank, 2018; Sarel, Finkelstein, Las, & Ulanovsky, 2017). These computational modeling papers demonstrate that a compressed ordered representation of future events is at least neurally plausible.

Previous work on neural correlates of statistical learning has shown that temporal proximity reflects in BOLD similarity and increases for predictive stimuli (Schapiro, Kustner, & Turk-Browne, 2012; Schapiro, Turk-Browne, Norman, & Botvinick, 2016; Turk-Browne, Scholl, Johnson, & Chun, 2010), particularly in the hippocampus (see Davachi and DuBrow (2015) for a review). In general, hippocampal BOLD activity is known to increase during sequence learning paradigms (Schendan, Searl, Melrose, & Stern, 2003; Ross, Brown, & Stern, 2009) and patients with bilateral hippocampal damage had impaired performance in learning sequential associations (Schapiro et al., 2012). The present study provides strong evidence that the representation of the future has a temporal organization, with logarith-mic compression. It remains to be seen whether this temporal organization can be captured with neuroimaging methods.

## Conclusion

This study introduced a novel Judgment of Imminence (JOI) task to study temporal order judgments for the future. The JOI task was designed to mirror the JOR task, allowing direct comparison (however, since JOI task required participants to learn a sequence of letters prior to testing, there were several methodological differences between the two tasks). The classic finding in JOR is that the response time (RT) for correct judgments varies as a function of the lag to the more recent probe and does not depend on the lag to the less recent probe. Based on this finding, many authors suggested a backward self-terminating search model operating on an ordered representation of the past. In the JOI task, it was found that correct RT depends on the lag to the more imminent probe and does not depend on the lag to the less imminent probe. This suggests a forward self-terminating search model operating on an organized representation of the future. These findings were supported by detailed consideration of RT distributions. This result supports the hypothesis that there is a deep symmetry between the psychological processes that underlie our ability to remember the past and make predictions about the future. Both memory for the past and predictions of the future appear to rely on a compressed, ordered representation.

## Acknowledgment

We thank Rebecca DiDomenica, Jodi Manning, Laxmi Nair, Arun Sidhu, Anita Wong, Sarah Dillard and Michelle Wiciak for their help in collecting the data.

## Methods

### Experiment 1: Judgment of Recency (JOR)

The procedure of this experiment followed closely the procedure of Experiment 2 of Hacker (1980). Participants were presented with a list of 9, 11, or 13 consonants at the rate of 5.5 letters per second. At the end of the list, two of the last seven letters were chosen randomly and the participants were asked to indicate using the left or right arrow key which of the two letters had appeared more recently. In Figure 1b, g and t are presented as the probe items. Because g was presented more recently than t, the correct answer is g. In addition, participants were asked to respond with the up arrow key if they did not remember seeing either of the probe letters on the list. If the participant did not make a response within 6 s, the trial was terminated. Less than .4% of trials terminated without a response. “UP” response was used in 17% of trials.

The distance to the more recent probe stimulus was varied from lag −1 (the last stimulus in the list) to −6. The lag of the less recent probe varied from −2 to −7. This leads to 21 possible combinations of lags, which were presented in random order. Each participant completed 320 trials.

There were several methodological differences between this procedure and the procedure of Experiment 2 of Hacker (1980). Unlike the Hacker (1980) study, in this experiment participants were never given foils that did not appear in the list. Also, in the Hacker (1980) study participants were not given the option to respond indicating that they did not remember either of the probes. The participants in the Hacker (1980) study were also more experienced in the task, experiencing a variety of presentation rates over several experimental sessions. In this experiment, participants received only one presentation rate in one session lasting about forty minutes.

#### Participants

The participants that participated in the study were drawn from the participant pool for Boston University’s introductory psychology class. The study materials and protocol were approved by the Institutional Review Board at Boston University. 108 participants signed up for the study. One participant withdrew from the experiment. Data from 11 participants were excluded because their overall accuracy was no better than chance.

### Experiment 1: Judgment of Imminence (JOI)

In order to probe the ability of participants to make relative order judgments of the future, participants were first trained on a probabilistic sequence. After training, the participants continued to see the same sequence interspersed with the probes consisting of two items that were from training. Participants indicated which of the two items would appear sooner (more imminent).

### Sequence generation

For each participant, a unique transition ring similar to the one shown in Figure 1c was generated by randomly selecting 13 consonants. Throughout the duration of the experiment, the participants saw letters drawn from this ring. With a probability of 0.9, the participants saw the next letter in the sequence. With probability 0.1, a random letter was chosen from the sequence that wasn’t the letter just shown to the participant. A presentation list of 1550 letters was generated using this procedure and presented to the participants one at a time over the course of the entire experiment from beginning to end. Each letter was displayed on the screen for 1.2 s followed by a fixation cross for .2 s.

### Training

During the training round, participants were instructed to repeat each letter silently to themselves and that they would be tested on this sequence. After sixty letters were shown to the participants (about 1.5 min), the initial training round ended and the experimenter initiated an abbreviated practice round for the actual task with the relative judgment probes (see below). The sequence continued for about two minutes. During this time participants were given twenty probes asking them to make predictions about relative order judgments of the future. The goal of this task practice round was to show the participants what the final task is so that they are motivated to pay attention to the sequence. After two minutes of this round, the training resumed and the participants were shown just the training sequence without any probes for another three minutes.

### Probes

After each letter from the list, a probe was shown with a probability of 0.3. The probe consisted of two letters that were presented side by side. The participants were told that they will continue to see the letters one at a time and that sometimes two letters will be presented side by side. The task was to predict which of the two letters will appear sooner using the “Left”/“Right” key. For instance, in Figure 1c, t and g are presented and the participant is asked to predict which of the two is more imminent. The correct answer here is t since it is two steps from the current position in the sequence *vs* g which is five steps in the sequence. The instructions clarified that they were not necessarily required to choose the next letter in the sequence (although that would sometimes be the case) but the one that would appear sooner. Probes were not shown one after another; at least one letter from the sequence was displayed before the next probe. If the participants failed to make a response in 4 s, they were shown a message,“Faster” for 0.2 s and then the task continued. About 1.3% of trials resulted in a time-out. On average the participants were shown 406 probes. The first 10% of the probes (these included the probes shown in training) were omitted from further analyses. The post-training round lasted for about thirty minutes with two intermediate breaks.

Based on pilot studies and debriefing interviews, it was found that participants were particularly cognizant of pairwise associations between the letters in the sequence. In order to eliminate an account based on simple pairwise associations between the probes, we did not present probes that contained stimuli that would be expected to be presented successively. For instance, in Figure 1c, if the more imminent item in the probe was x, the less imminent item could not be q. The two probe letters were sampled from the next 7 positions in the transition ring. The lag of the more imminent item varied from 1 (next in the sequence) to 5. For each of these more imminent items, the less imminent items were chosen using predetermined combinations. The points were sampled such that, where possible, there were three less imminent probes for each more imminent probe. For instance, when the lag of the more imminent probe was 1 (next item), the less imminent item was chosen from a lag of 3, 5, or 7. When the more imminent probe was at a lag of 3, the less imminent probe was chosen from lags of 5, 6, or 7. Since only the next 7 positions are probed, a more imminent probe at a lag of 4 can only be paired with a less imminent probe at a lag of 6 and 7. A more imminent probe at a lag of 6 can only be probed with a more imminent pairing at a lag of 7.

### Participants

Sixty healthy young adults were recruited from Boston University to participate in one session each and were paid $15 per hour for their time. The study materials and protocol were approved by the Institutional Review Board at Boston University. Three participants performed no better than chance; their data were excluded from further analysis.

### Experiment 2: Same participants performing JOR and JOI

The experimental procedure closely followed the procedure of Experiment 1. 32 participants performed both JOR and JOI (it was chosen randomly whether a participant will do JOR or JOI first). Each participant completed both tasks in the same day.

Overall, the participants performed similarly to Experiment 1. In JOR they did not make a response in about 0.5% of trials and they selected “UP” response in about 12.6% of trials. In JOI the participants did not make a response in about 2.3% of trials. In JOR data from 1 participant and in JOI data from 9 participants were excluded because their overall accuracy was no better than chance. Overall, 22 participants performed both tasks above chance.

### Strategy for analyzing the two lag variables

On each trial the participant is given two probe stimuli. We are interested to know if the lag of both of the probes affects correct RT or if only the lag of the correct probe does. One might have simply put these two variables (perhaps recording them using a distance between the two probes) into a linear regression. However the recencies/imminencies are not sampled equally. Moreover, the effect of lag appears to be non-linear (Fig. S3). Using a straightforward linear model carries the risk of a spurious distance effect due to residuals from a non-linear and unequally sampled relationship.

In order to control for these effects, the following two-step strategy was used. First, the distance effect was quantified by calculating the slopes of the lines joining the dependent variable as a function of the less salient independent variable. For instance, in Figure S2 the lines for the correct RT appear to be flat. Thus the median RT does not appear to depend on the lag of the less recent probe. In order to quantify this intuition, the slope of each line was calculated for each participant. In order to ascertain whether these slopes are meaningfully different from 0 or not, a Bayesian t-test (Rouder et al., 2009) was performed on the slopes obtained for each participant. If a particular variable did not contribute to the dependent variable, it was excluded as a factor in a subsequent linear mixed effect analysis. Thus the purpose of this analysis was simply to ascertain whether a factor meaningfully contributed to the dependent variable. Once the relevant factors affecting the dependent variable were ascertained, further analyses were performed with the relevant variables. The function of the linear mixed effect analysis is simply to determine whether the apparent effect of the more salient variable is statistically reliable. Error bars in Figures S1, S2, S3, S4 and S6 represent the 95% confidence interval of the mean across participants normalized using the method described in (Morey, 2008).

## Supplemental Information

As supplemental information, we provide details of several additional analyses. Specifically, we show the data supporting the conclusion that the accuracy was a function of a lag of both more and less recent/imminent probe item. For JOR we show that incorrect responses deepened only on the lag of the selected probe. For JOI the results of the analysis of incorrect responses did not lead to strong evidence. Here we also provide broken out RT plots for each of the two experiments (figures in the main text included data from both experiments put together). We also show details of the analysis that compares performance in JOR and JOI in Experiment 2 (in Experiment 2 the same subjects did both tasks). We did not find a significant correlation for either RT, accuracy, or scanning rate.

**Figure S1.**
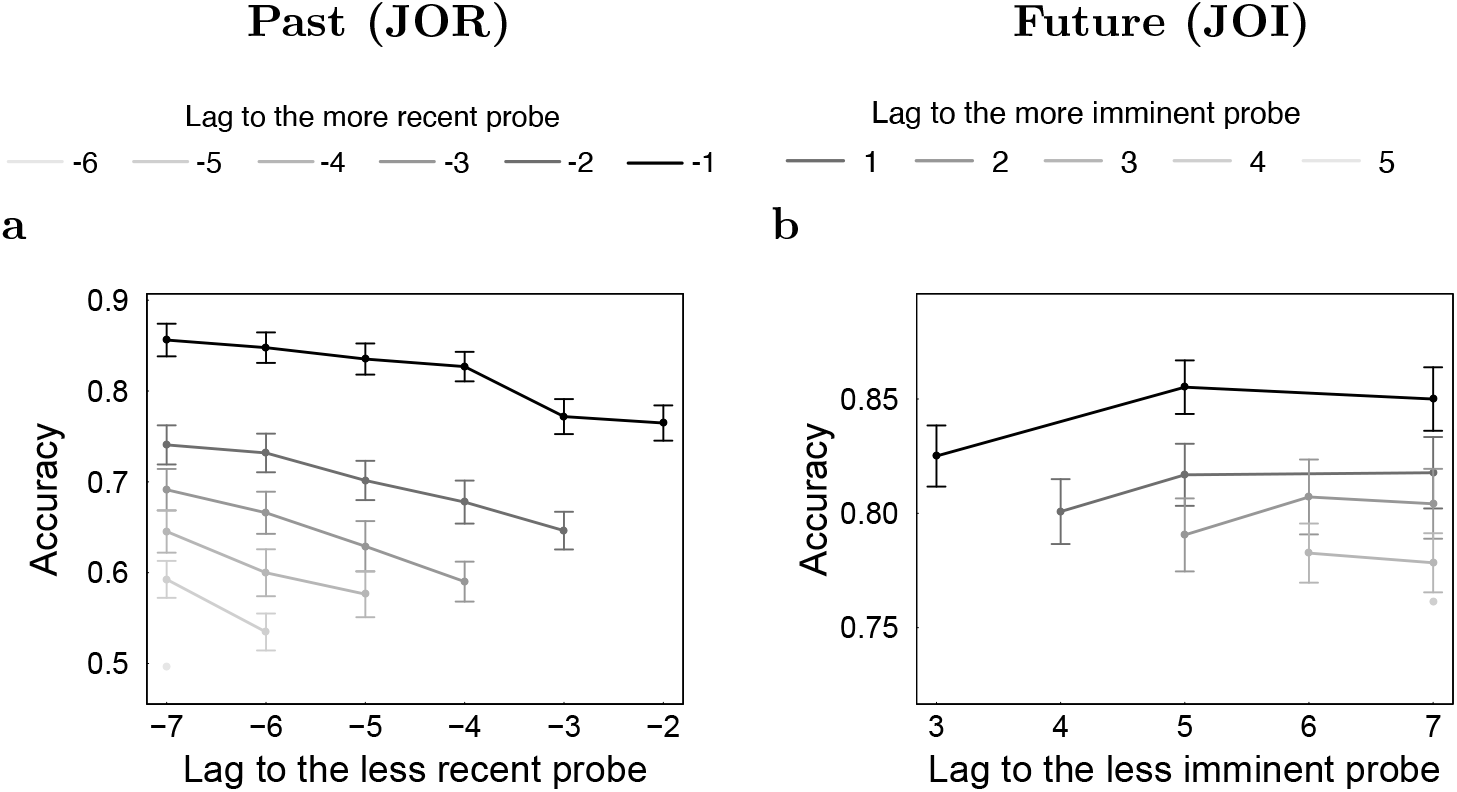
Accuracy in JOR and JOI. **a.** Accuracy in JOR showed a reliable recency effect and a reliable distance effect. **b.** Accuracy in JOI was higher for probes that were closer in time to the present but did not show a reliable distance effect. See text for details.

### JOR accuracy showed a recency effect and a distance effect

The probability that participants selected the more recent probe was .70 ± 0.01 in Experiment 1 and .69±0.01 in Experiment 2 (mean accuracy across participants, taking into account only trials when participants responded by selecting one of the probes - excluding “UP” responses and trials that ended with no response). The accuracy was .82 ± .01 when the lag of the more recent probe was −1 and dropped to .49 ± .02 when the lag was −6 in Experiment 1. Similarly, in Experiment 2, the accuracy was .81 ± .02 when the lag of the more recent probe was −1 and dropped to .51 ± .03 when the lag was −6. In Experiment 1 at lag −6 the probability of choosing the more recent probe was not different from chance (Chi-squared prop test, *χ*^2^(96) = 89.1, p-value not significant). Lag −5 had an accuracy of 0.56 ± 0.01 and was significantly higher than chance (Chi-squared prop test, *χ*^2^(96) = 142.6, *p* < 0.01). In Experiment 2 at lag −6 and −5 the probability of choosing the more recent probe was not different from chance (for lag −6: Chi-squared prop test, *χ*^2^(31) = 29.34, p-value not significant; for lag −5: Chi-squared prop test, *χ*^2^(31) = 44.28, p-value not significant). Lag −4 had an accuracy of 0.60 ± 0.02 and was significantly higher than chance (Chi-squared prop test, *χ* (31) = 90.52, *p* < 0.01).

**Figure S2.**
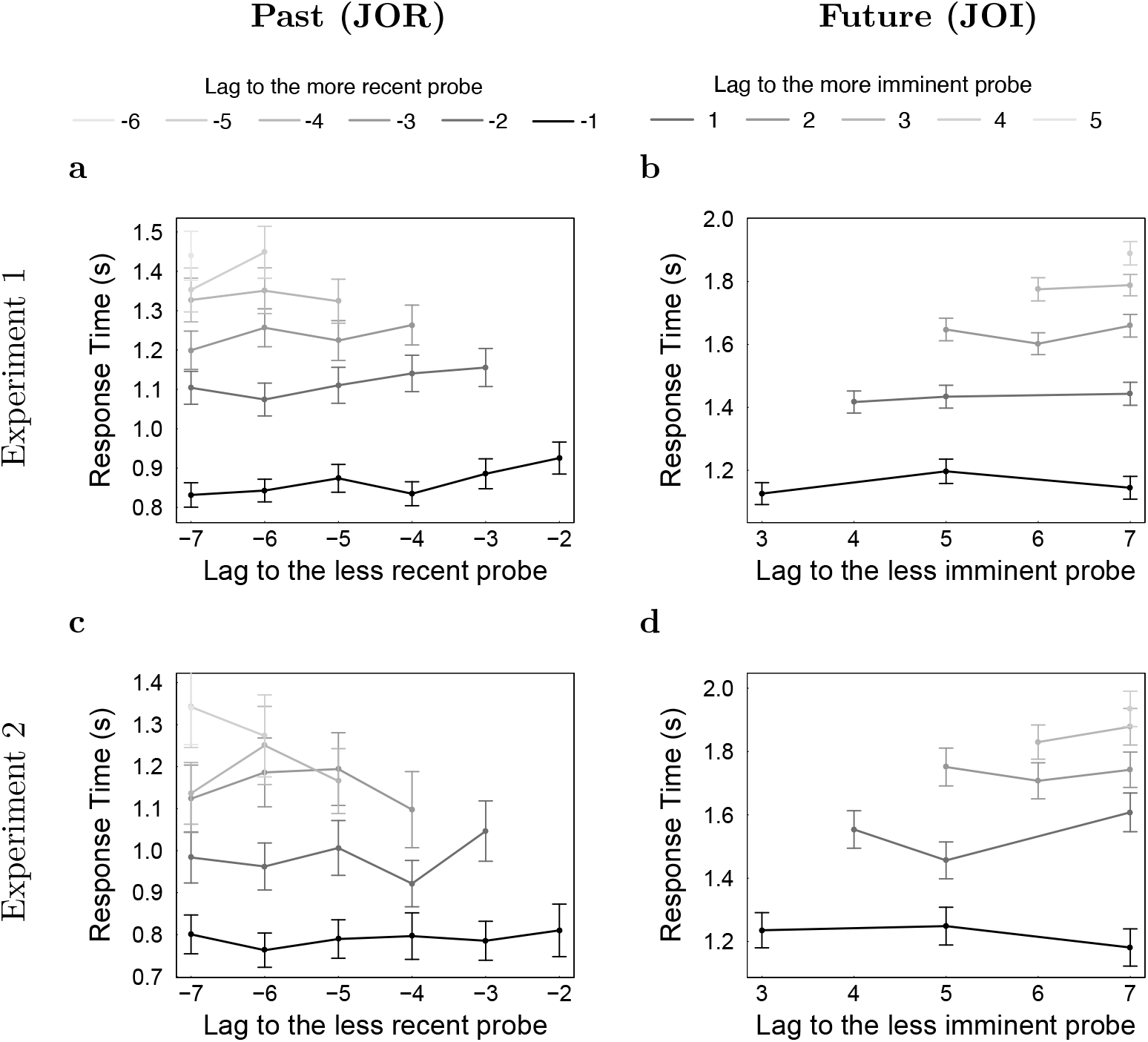
Analogous to Figures 3a and b but with results for each experiment shown separately. Median correct RTs depend on the lag of the more recent/imminent probe but not on the lag of the less recent/imminent probe. This is consistent with a serial self-terminating search model.

Accuracy also depended on the temporal distance between the more recent probe and the less recent probe. For a lag of the more recent probe, the accuracy improved as the lag of the less recent probe decreased (distance effect). The upward-sloping lines in Figures S4a and c indicate the presence of this distance effect. To quantify the distance effect for each participant, we calculated the slope of each line in Figure S4a and c. A Bayesian t-test (Rouder et al., 2009) on the obtained slopes revealed “decisive evidence” (Wetzels & Wagenmakers, 2012; Kass & Raftery, 1995; Jeffreys, 1998) favoring the hypothesis that the slopes are different from 0 (JZS Bayes Factor > 10^2^ in both Experiment 1 and Experiment 2).

The effects of lags of the two probes on accuracy was quantified using a linear mixed effect analysis with independent intercepts for each participant. The accuracy increased with an increase in the lag of the more recent probe by .077 ± .002, *t*(1918) = −31.9, *p* < 0.01 per unit change in lag in Experiment 1 and by .070 ± .004, *t*(618) = −16.49, *p* < 0.01 per unit change in Experiment 2. Accuracy also increased with the lag of the less recent probe by .023 ± .002, *t*(1918) = 9.73, *p* < 0.01 per unit in Experiment 1 and .028 ± .004, *t*(618) = 6.56, *p* < 0.01 per unit in Experiment 2. These findings are consistent with the findings from prior studies.

**Figure S3.**
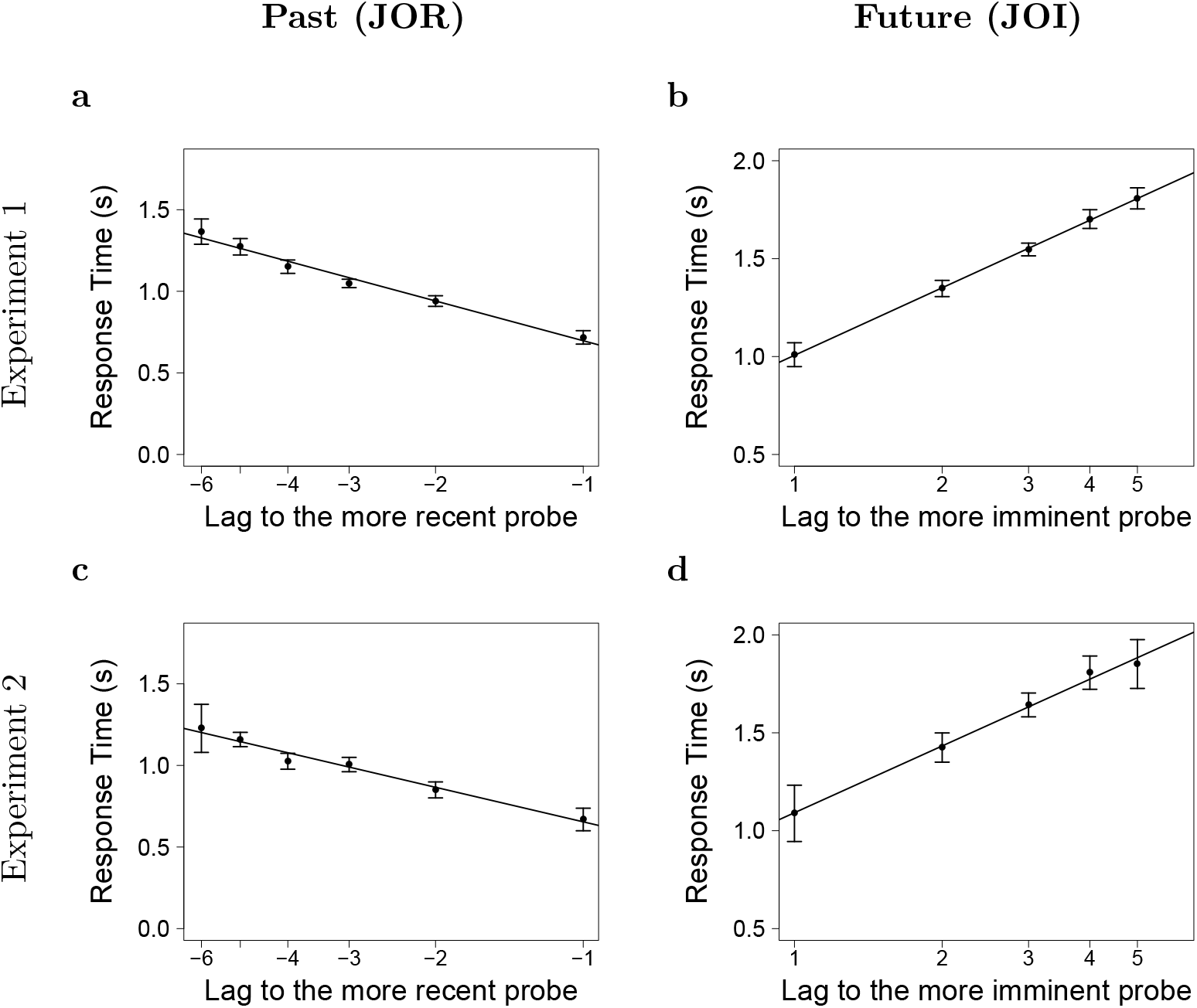
In each of the two experiments median correct RT varied sub-linearly with lag. This figure is analogous to Figure 3c,d, but with results for each experiment shown separately.

In JOR we observed that less recent item had an impact on accuracy, but not on RT, causing the distance effect only on accuracy. To explain this in the context of serial scanning mechanisms we discuss two possible causes for participants to make an incorrect judgment according to scanning models. First, participants might not properly encode the more recent probe and simply scan past it until reaching the less recent probe. In that case, the lag of a less recent probe does not impact the accuracy. Second, the noise level in the evidence accumulation process might be large enough such that participants do not accumulate enough evidence for the more recent probe when they reach the less recent probe during the scanning process. Consistent with the second explanation, the results reported here support a hypothesis that items on the timeline are represented with broad overlapping fields that have sequential peaks.

**Figure S4.**
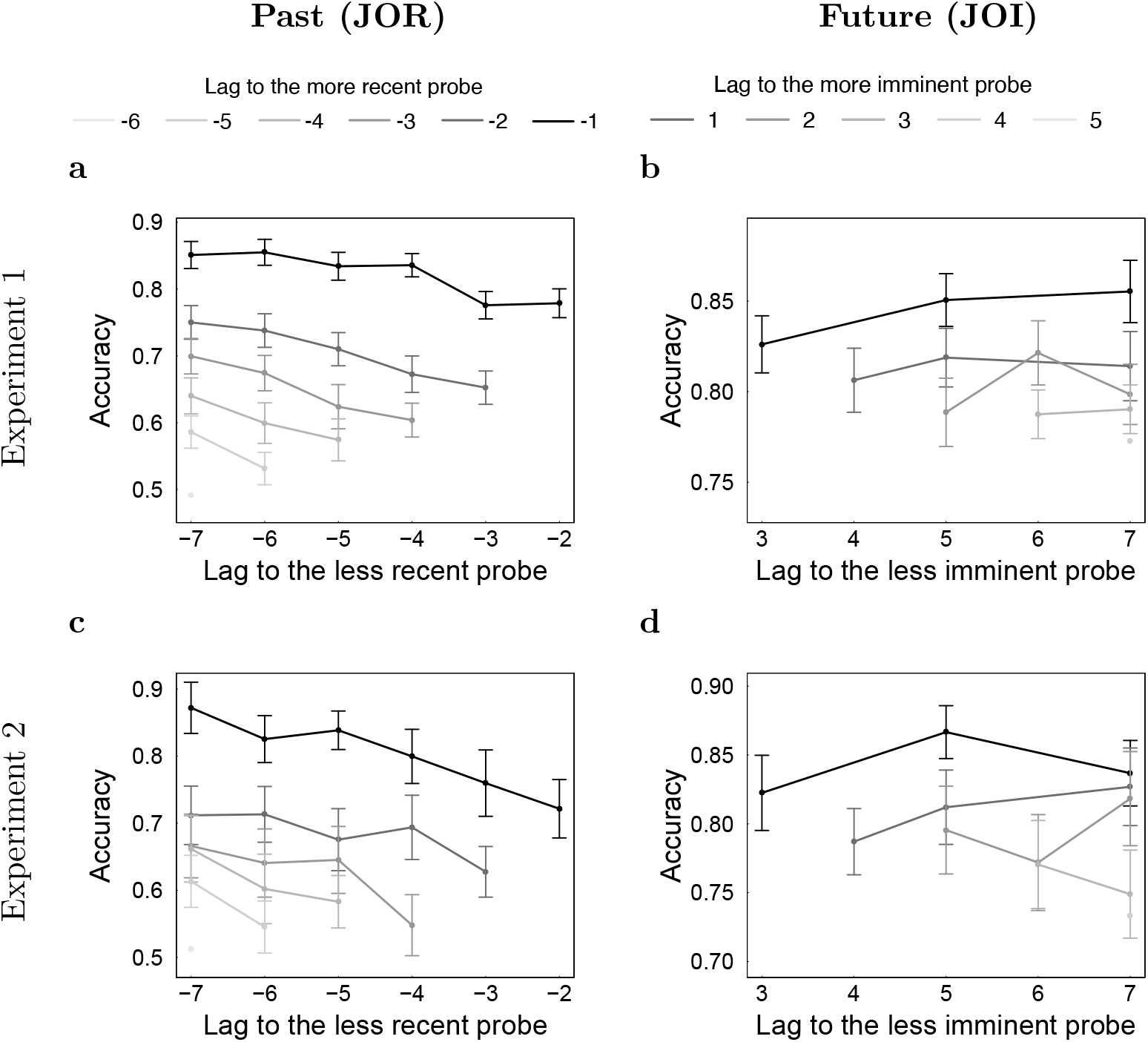
Accuracy in JOR and JOI across the two experiments. This figure is analogous to Figure S1, but with results for each experiment shown separately. In both experiments, accuracy in JOR showed a reliable recency effect and a reliable distance effect. Also, in both experiments accuracy in JOI was higher for probes that were closer in time to the present but did not show a reliable distance effect.

### Incorrect RT depended only on the lag of the selected probe in the JOR task

In a self-terminating backward scanning model, if the scan misses the more recent probe, it would then terminate on the less recent probe. These responses would be errors and the scanning time for these errors would depend on the lag to the less recent probe.

Given the overall error rate of .30 ± 0.01 in Experiment 1 and .31 ± 0.01 in Experiment 2, there was less than half the number of observations for incorrect RTs as there was for correct RTs. Also note that the number of errors was not evenly distributed over lags, so that some points have many fewer observations than others (Figure S5). Nonetheless, error RTs appear to depend reliably on the lag to the less recent item. There does not appear to be a strong effect of the lag to the more recent probe.

**Figure S5.**
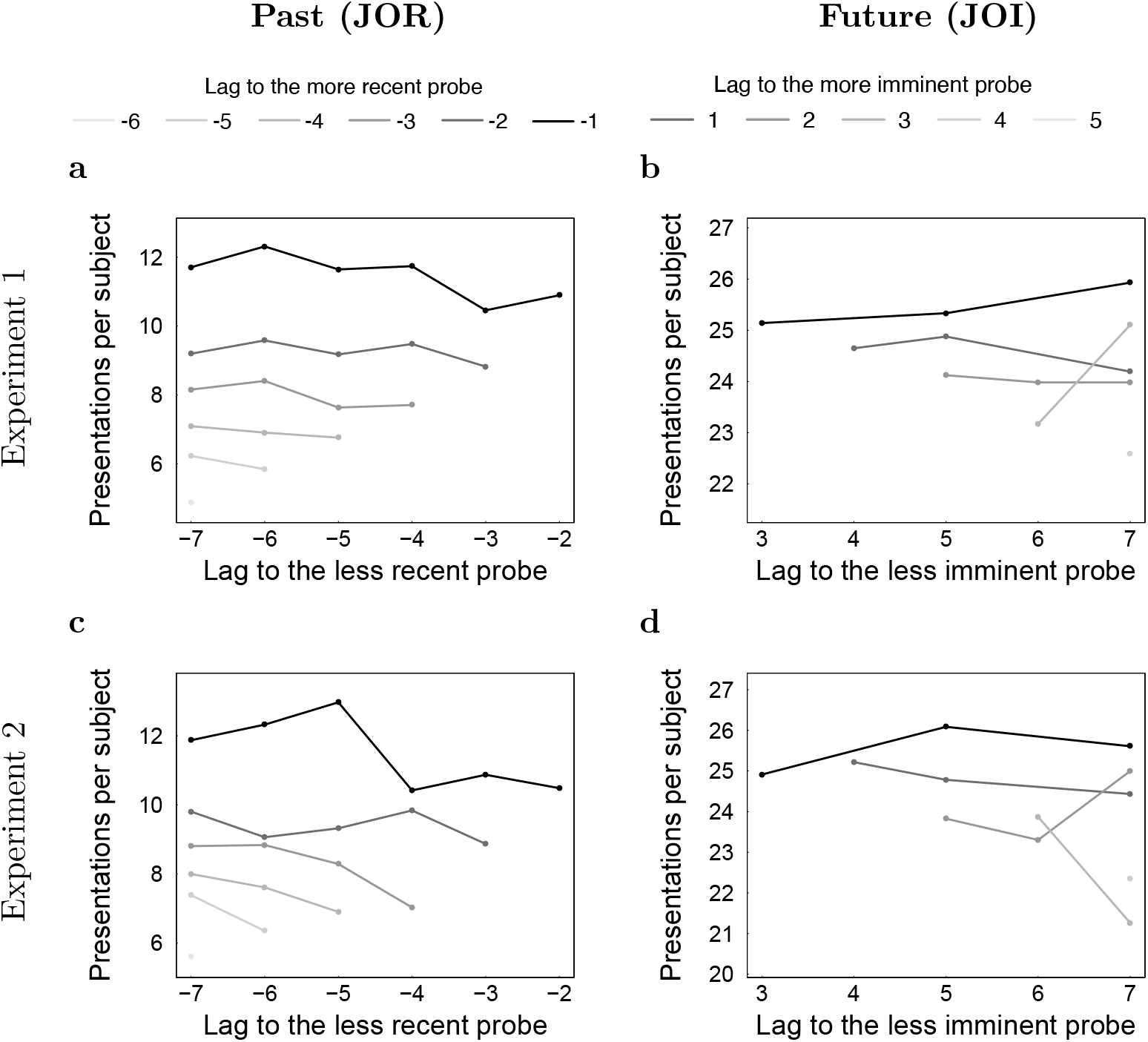
Number of probe presentations for every available combination of more and less recent/imminent lag in correct trials. Since in JOR participants had an option to press “UP” button to indicate that they do not remember seeing either of the probes, the number of correct presentations was lower in JOR compared to JOI.

To evaluate whether there was an effect of the lag to the more recent probe we calculated the slope of the distance effect for each value of the lag to the less recent probe. This is analogous to the distance effect calculation for correct RTs except the distance effect is calculated separately for the lag to the less recent probe rather than the more recent probe. A Bayesian t-test showed “strong evidence” favoring the hypothesis that the slopes of the median RTs as a function of the more recent lag are not different from 0 (JZS Bayes Factor = 14.5) in Experiment 1 and “substantial evidence” (JZS Bayes Factor = 5.79) in Experiment 2.

A linear mixed effects analysis, allowing each participant to have an independent intercept, and the less recent lag as regressor showed a significant effect of the lag to the less recent probe on the median RT of incorrect responses in Experiment 1 (.033 ± .007 s, *t*(473) = 4.9, *p* < 0.001), but not in Experiment 2 (.019 ± .012 s, *t*(153) = 1.57).

**Figure S6.**
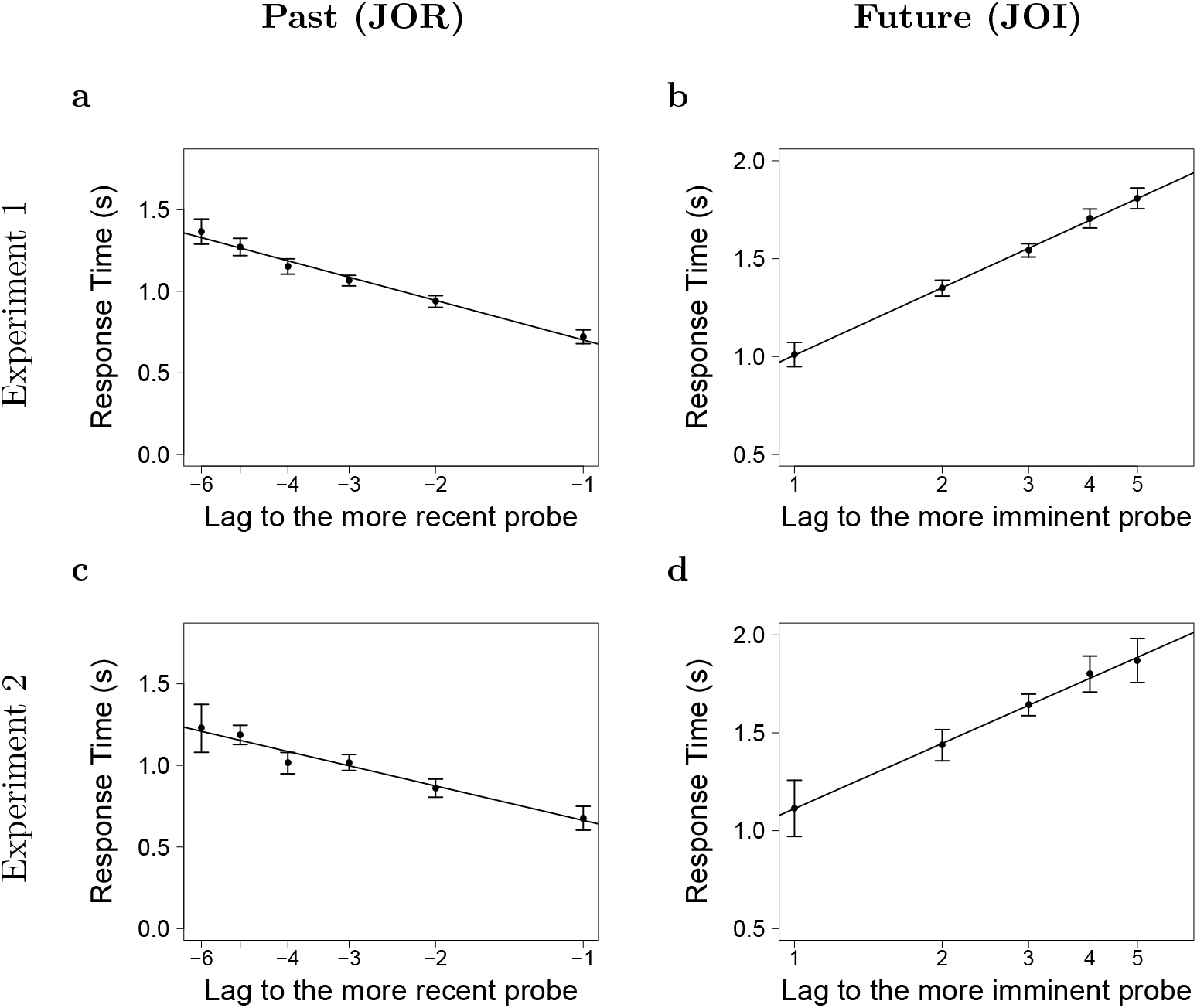
Sublinear relationship between lag and RT was preserved when the dataset was modified such that each pair of more and less recent/imminent lag had the same number of correct trials (the number of observations for each pair was reduced to the number of observations of a least frequent pair by performing random sampling).

### JOI accuracy depended only on the lag of the more imminent probe

The overall accuracy in JOI was .81 ± .02 in Experiment 1 and .80 ± .03 in Experiment 2. In Experiment 1 the accuracy varied from .84 ± .02 when the most imminent lag was 1 to .77 ± .02 when the most imminent lag was 5. In Experiment 2 the accuracy varied from .84 ± .03 when the most imminent lag was 1 to .73 ± .02 when the most imminent lag was 5.

In order to check if the somewhat upward-sloping lines in Figures S4b and d indicate the presence of a reliable distance effect, the slopes of each of the lines were calculated for every participant. A Bayesian t-test (Rouder et al., 2009) on the obtained slopes revealed “substantial evidence” favoring the hypothesis that the slopes are not different from 0 (JZS Bayes Factor 7.2 in Experiment 1 and 8.53 in Experiment 2).

Thus the accuracy depended on the lag to the more imminent probe alone. The effects of the more imminent probe on accuracy were quantified using a linear mixed analysis with independent intercepts for each participant. Accuracy decreased with an increase in the lag to the more imminent probe by .017 ± .002, *t*(227) = −6:9, *p* < 0:01 per unit change in lag in Experiment 1 and by .026 ± .004, *t*(91) = −6:9, *p* < 0:01 in Experiment 2. The finding that participants were more accurate based on the lag of the more imminent probe is different from the accuracy results seen in JOR (Hacker, 1980) where accuracy depends on both the more recent and the less recent probes. Keeping in mind that JOI had fewer pairs of more and less imminent lags than JOR, an additional reason for this finding could be in the fact that in JOR participants could use “UP” button to indicate that they do not remember seeing the probe items in a given trial. Availability of “UP” button in JOR and not JOI was motivated by the fact that in JOI participants have seen all the items many times. On the other hand, if participants did not pay much attention during a particular JOR trial, they might not remember seeing the probe items. However, this also resulted in participants making more guesses in JOI than in JOR, making it harder to find the distance effect.

**Figure S7.**
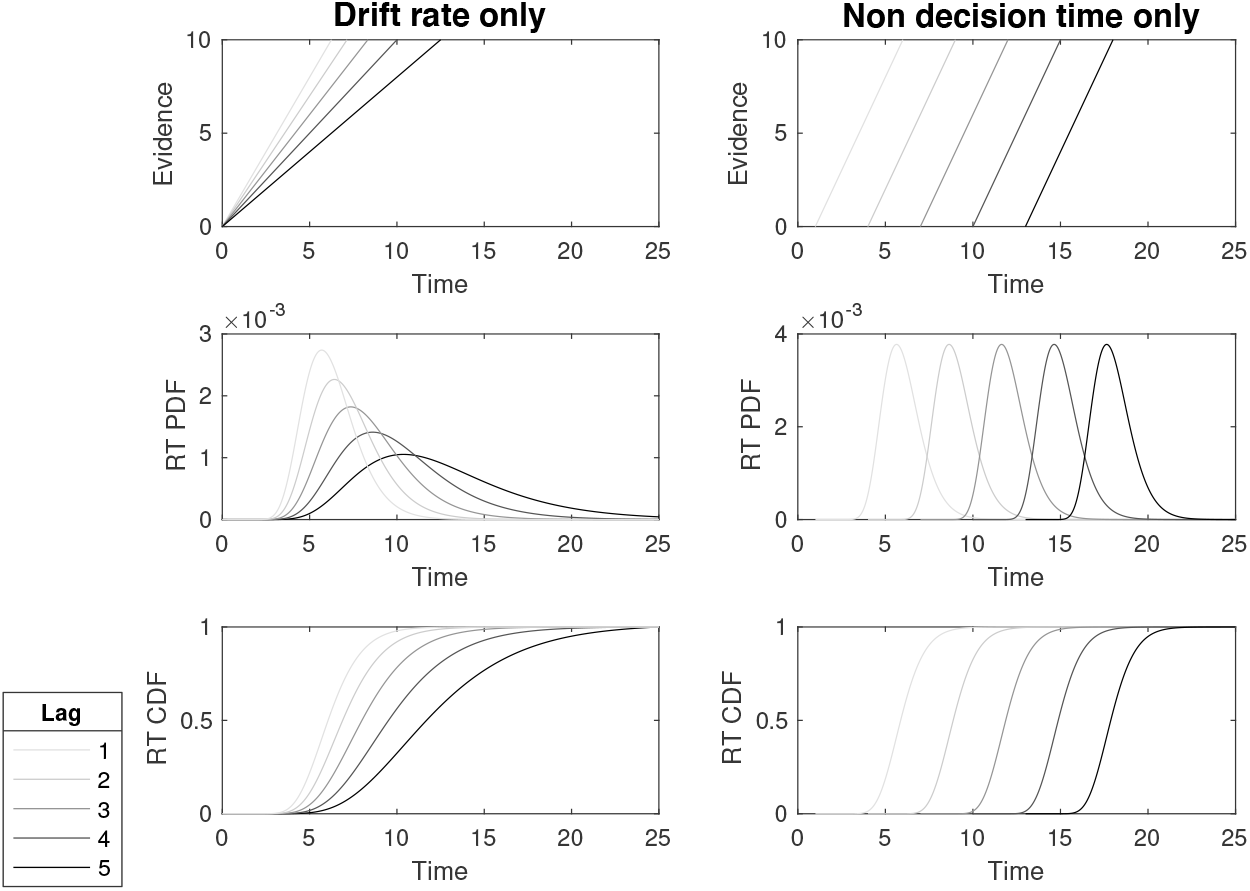
Illustration of two memory search hypotheses: parallel and serial. In parallel models evidence for each item is immediately available and it is accumulating at a larger rate for items with smaller lag (top row). Therefore the RT distributions for different lags have the same origin, but they increase at different rates (middle and bottom row). In serial models items are accessed in sequential order, therefore the evidence for items with smaller lags starts to accumulate before the evidence for the items with larger lags (top row). Consequently, RT distributions for different lags are shifted with respect to one another (middle and bottom row).

### Incorrect RT showed mixed results in the JOI task

When participants make errors, the forward scanning model predicts that the RT should depend on the lag of the less imminent probe. However this was not observed in the error responses here, perhaps because accuracy was much higher than in JOR. In Experiment 1 the overall error rate was .19 ± .02 and for each participant on average 6 error responses per condition were obtained. In Experiment 2 the overall error rate was .20 ± .03 and for each participant on average 3 error responses per condition were obtained.

**Figure S8.**
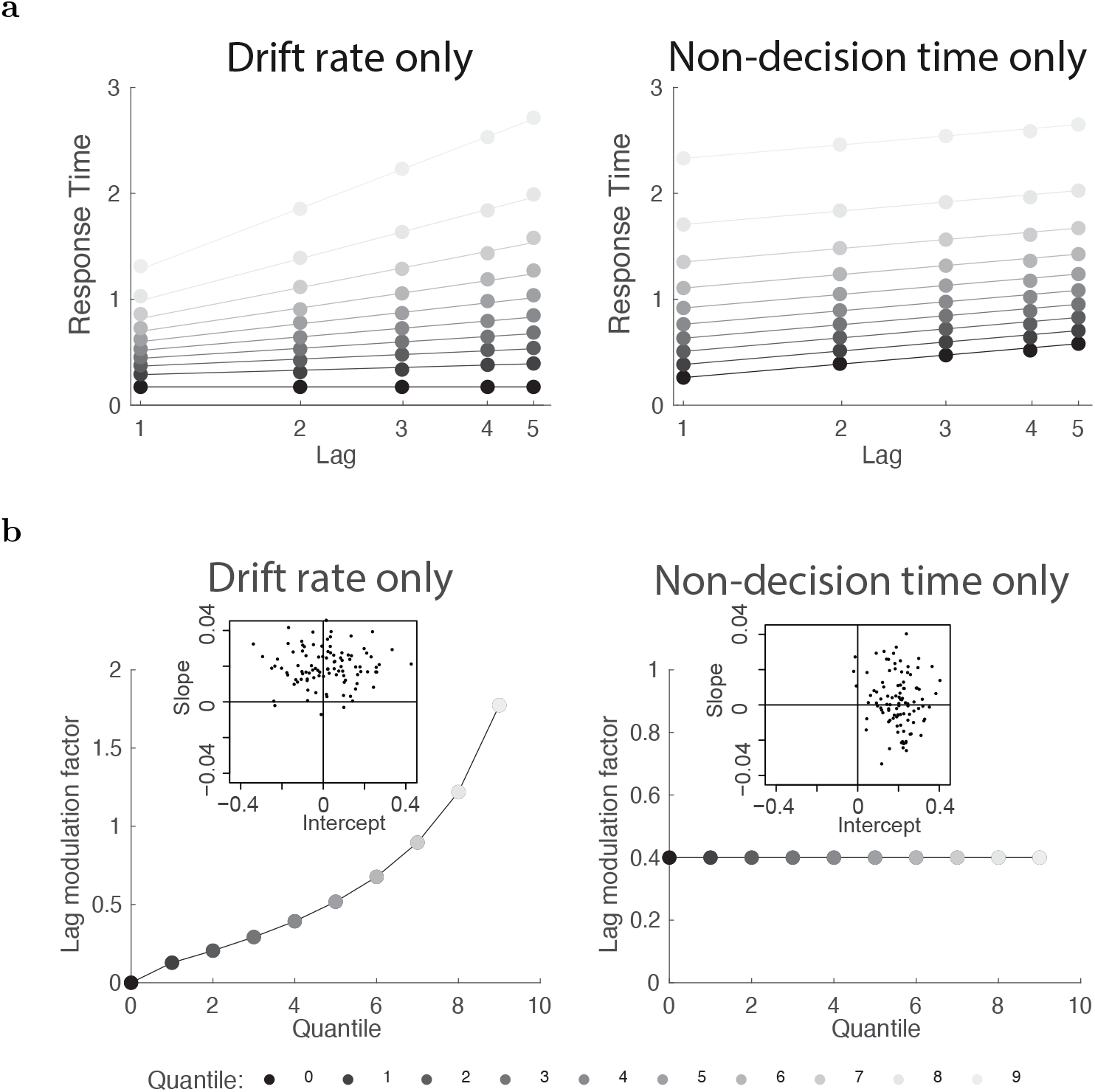
Properties of RT distributions according to “drift rate only” (consistent with parallel search models) and “non-decision time only” (consistent with serial search models) hypotheses. **a.** Deciles of the RT distributions as a function of log lag are distinct for the two hypotheses. In parallel models evidence for each item is immediately available and it is accumulating at a larger rate for items with smaller lag. Therefore the RT distributions for different lags have the same origin, but they increase at different rates. As a consequence, the parallel search hypothesis predicts that the effect of lag should decrease with the quantile of the RT distribution. In serial models, items are accessed in sequential order. Therefore the evidence for items with smaller lags starts to accumulate earlier than the evidence for the items with larger lags. Consequently, RT distributions for different lags are shifted with respect to one another. Thus, the serial scanning model predicts that quantiles as a function of lag should give parallel lines. **b.** Lag Modulation Factor is the slope of the lines in panel (a) for each quantile. The parallel model predicts that the Lag Modulation Factor should decrease systematically with quantile and should asymptotically be zero. The serial search model predicts that the Lag Modulation Factor should not be affected by quantile and should asymptote at some non-zero value. The inserts illustrate hypothetical results across participants for each hypothesis, with the slope and intercept of the Lag Modulation Factor as a function of quantile.

**Figure S9.**
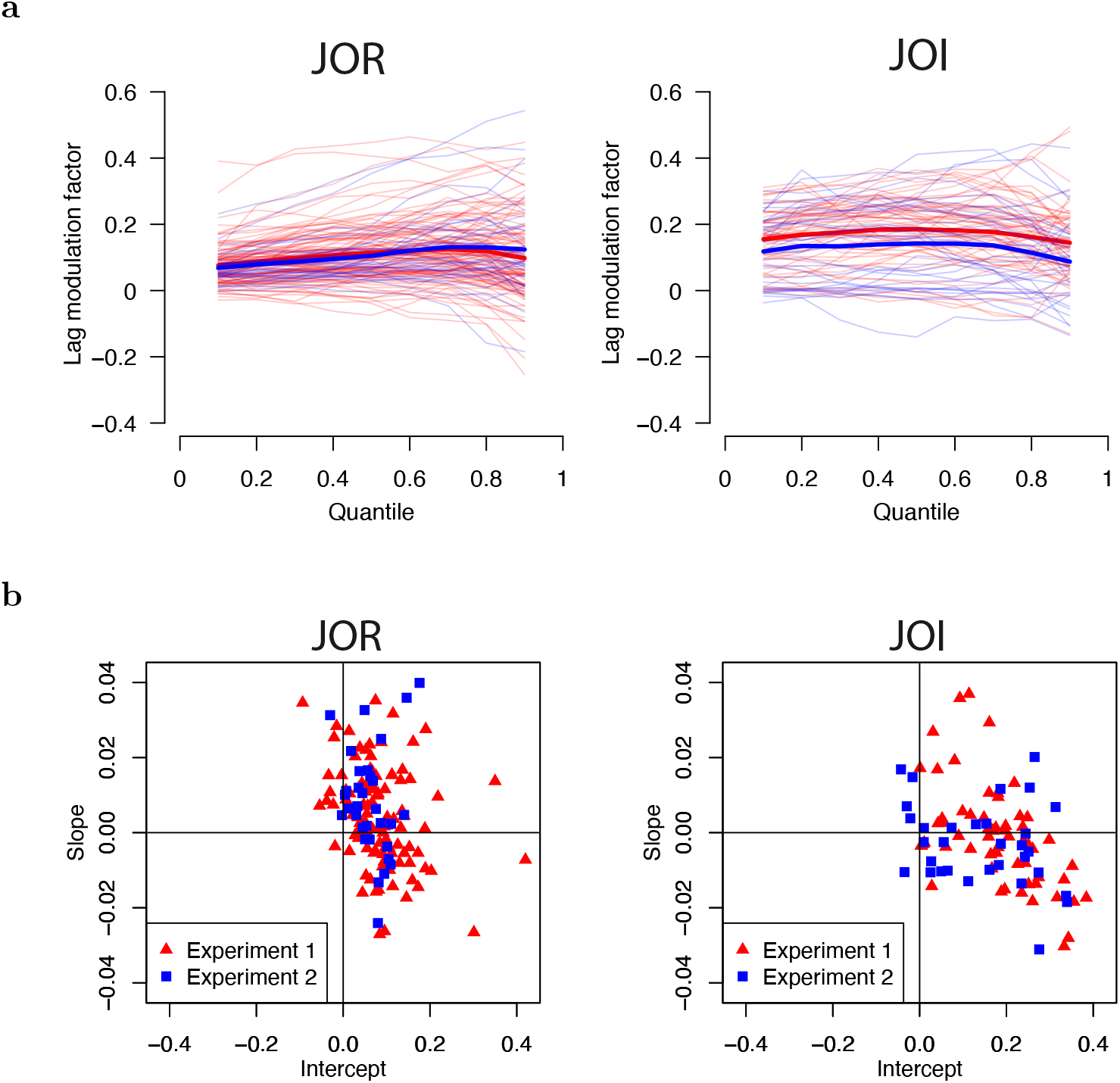
Properties of RT distributions were consistent with the scanning hypothesis in both JOR and JOI. **a.** Lag Modulation Factor as a function of quantile for each participant. Participants in Experiment 1 are shown with light-red lines and in Experiment 2 with light-blue lines. The mean across participants for each experiment is shown with a thick line of the appropriate color. **b.** Slopes and intercepts of the Lag Modulation Factor for all the participants. For both JOR and JOI the results appear more similar to those predicted by the serial model than to those predicted by the parallel model.

**Figure S10.**
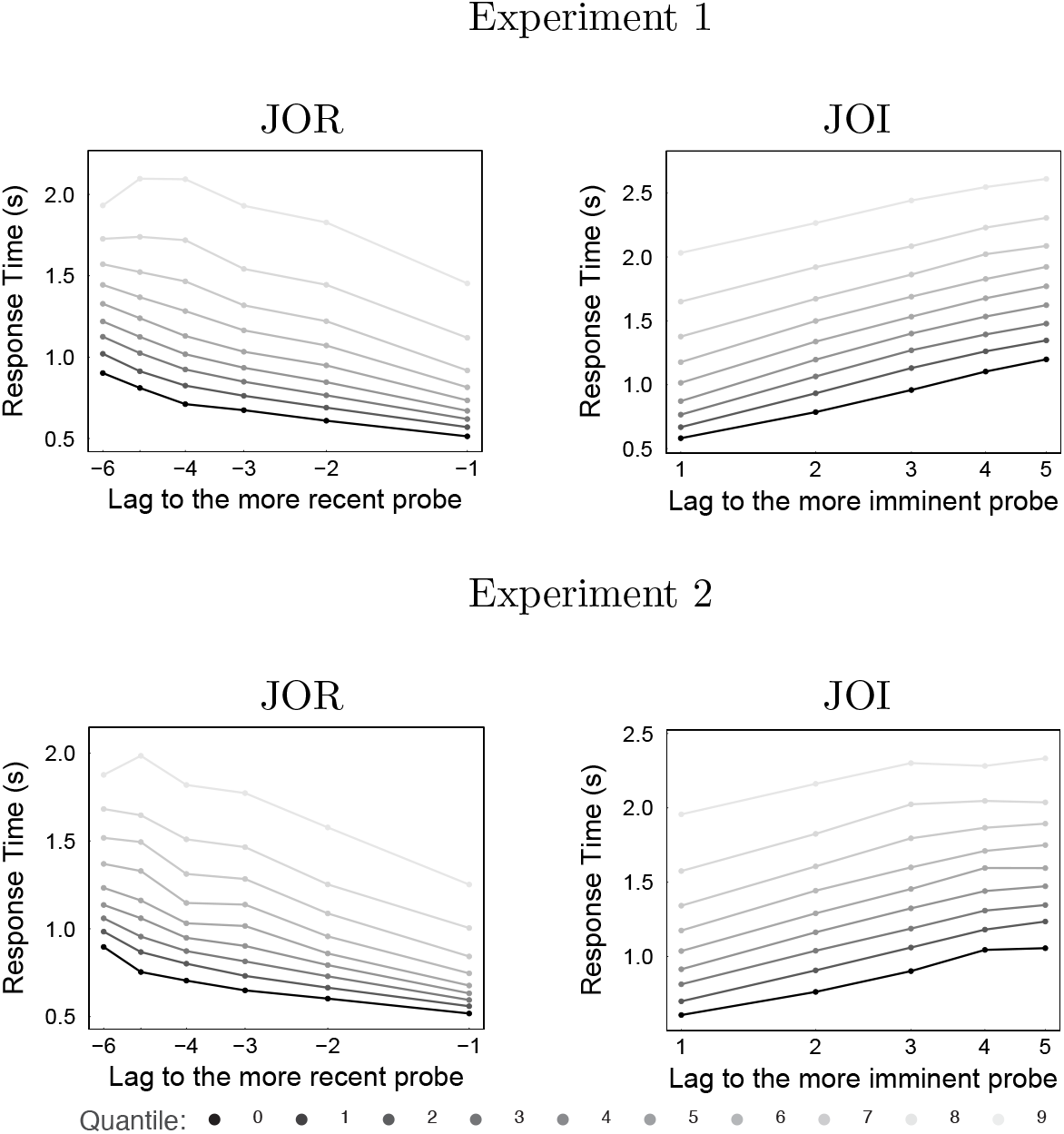
Response time as a function of lag for every decile across all the participants separated per experiment. The results were consistent across the two experiments. See Figure 4 and the main text for more details.

To evaluate whether the more imminent probe contributed to the change in median RTs of a particular less imminent probe (distance effect), the slope across the more imminent lag for each less imminent lag was calculated. This is analogous to the distance effect calculation for correct RTs except the distance effect is calculated separately for the lag of the less imminent probe rather than the more imminent probe. The less imminent lags of 3 and 4 only have one possible combination so a slope could not be calculated for these lags. Across the remaining slopes, a Bayesian t-test showed that the evidence for a distance effect was “barely worth a mention” in both Experiment 1 and Experiment 2 (JZS Bayes Factor = 1.1 in both experiments).

Given the overall error rates of .19 and .20, there was less than one fourth the number of observations for incorrect RTs as there was for correct RTs. While a linear mixed effects analysis, allowing each participant to have an independent intercept, and both lag as regressors, showed a significant effect of the lag of the more imminent probe on the median RT of incorrect responses, .05 ± .05 s, *t*(586) = 2.9, *p* < 0.01 in Experiment 1 and .05 ± .03 s, *t*(228) = 2.1, *p* < 0.05 in Experiment 2, a linear model with both the more and the less imminent lag did not show any significant effect of the more or the less imminent probe, .04 ± .02 s, *t*(641) = 1.7 and .01 ± .02 s, *t*(641) = .4 respectively in Experiment 1 and .06 ± .04 s, *t*(250) = 1.4 and .00 ± .04 s, *t*(250) = .0 in Experiment 2.

### Comparison of cross-participant performance in JOR and JOI

Out of the 32 participants that performed JOR and JOI in Experiment 2, 22 were above chance in both tasks. We investigated the correlation of different measures related to response time and accuracy in JOR and JOI across those 22 participants. Briefly, we did not find reliable correlations in any of the dependent measures we considered. RT in JOR was not correlated with RT in JOI, *r*(20) = .41, *p* = .050. Similarly, accuracy was not correlated across participants in JOR and JOI, *r*(20) = .18. We also computed the slope of RT as a function of log lag of a more recent/imminent probe for each participant and found no correlation between JOR and JOI, *r*(20) = .01.

### Individual participant results

Additional supplementary material shows the results for each participant in each of the two experiments. For participants in Experiment 2, results from JOR and JOI are shown side by side for easy comparison.

## Footnotes

1 Although previous results are consistent with a sublinear increase in RT with lag (see e.g., Fig. 7 of Hockley, 1984, Fig. 5 Hacker, 1980) this was not statistically evaluated in those prior studies.

2 The same conclusions were reached using polynomial regression.

